# Complementation of a temperature sensitive *Escherichia coli rpoD* mutation using *Lactobacillus* sigma factors

**DOI:** 10.1101/003152

**Authors:** James D. Winkler, Katy C. Kao

## Abstract

Housekeeping sigma factors in the *σ*^70^ family, as components of the RNA polymerase holoenzyme, are responsible for regulating transcription of genes related to vegetative growth. While these factors are well understood in model organisms such as *Escherichia coli* and *Bacillus subtilis*, little experimental work has focused on the sigma factors in members of the *Lactobacillus* genus such as *Lactobacillus brevis* and *Lactobacillus plantarum*. This study evaluates the ability of putative *σ*^70^ proteins from *L. brevis* (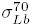) and *L. plantarum* (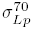) to complement a temperature sensitive mutation in the *E. coli* 285c *σ*^70^ protein. This report is the first to show that these heterologous sigma factors were capable of restoring the viability of *E. coli* 285c for growth at 40-43.5 °*C*, indicating the 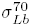 and 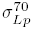 are capable of initiating transcription in a complex with the *E. coli* 285c RNA polymerase. These heterologous sigma factors may therefore be useful for improving biochemical knowledge of the sigma factor family or for use in the expression of hetereologous genomic libraries.

## Introduction

Two members of the *Lactobacillus* genus, *Lactobacillus brevis* and *Lactobacillus plantarum*, have been shown to possess many desirable characteristics including superior acid, butanol, and ethanol tolerance that are of interest for industrial usage [1–5]. These species have only recently been analyzed [6–9] and as such their biochemical and genetic corpus is lacking compared to the model organism *Escherichia coli*. In particular, the transcriptional programs of the *Lactobacilli* and regulation therein under many conditions are poorly understood, though characterization efforts are underway [10–12]. Housekeeping sigma factors (*σ*^70^ or *σ*^*A*^) [13] such as those putatively found in *L. brevis* and *L. plantarum* genomes, given their role in controlling gene transcription through promoter recognition during exponential growth, play an important role in the physiology of these organisms. Experimental study of the *Lactobacilli* sigma factors will improve our understanding of how these factors control the *Lactobacillus* transcriptional program.

The importance of sigma factors arises from their transient contact with the eubacterial RNA polymerase core enzyme (*α*_2_*ββ*′*ω*) that allows RNAP to specifically recognize promoter sequences upstream of genes [14–18]. Without these proteins, gene transcription is impossible. Sigma factors are divided into protein families based on structural homology, with factors responsible for regulating gene expression during exponential growth belonging to group 1 [13, 15, 19, 20]; by acting in concert, the various sigma factors within an organism effectively divide the transcriptional program of an organism into multiple sub-routines capable of responding to environmental conditions rapidly [21]. While the understanding of the biological roles of sigma factors found in well characterized organisms is extensive [22–26], the lack of experimental data on *L. brevis* and *L. plantarum* physiology means that the putative sigma factors in these organisms are understood only by analogy to the related *Bacillus subtilis* [27] and through sequence analysis of known promoters. In light of their role in regulating genes critical for growth, the housekeeping sigma factors of *L. brevis* (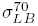) and *L. plantarum* (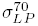) are the best candidates for *in vivo* examination to better understand the transcriptional programs of these organisms.

Analysis of the *Lactobacilli* sigma factors poses significant challenges given the relative paucity of genetic tools available for these organisms. We chose to examine the behavior of 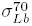 and 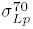 within *E. coli* as a result, in spite of the significant differences between the organisms. This approach has been successful in the past with non-*E. coli* housekeeping sigma factors [28–32] and other factors regulating flagellar production (*σ*^*F*^) and response to extracellular stress (*σ*^*E*^) [33–35] in complementing *E. coli* null phenotypes. Sigma factors that are too structurally divergent from those found in *E. coli* tend to be non-functional *in vivo* [36]. Given the diversity of heterologous sigma factors that are functional in *E. coli*, there must exist a significant amount of functional flexibility of *σ*-RNA polymerase interactions. An improved understanding of the minimal sigma factor sequence required for functioning would improve our understanding of both a basic aspect of bacterial physiology and enhance efforts for sigma factor engineering.

To that end, this work analyzes the functionality of 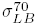 and 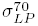 within an *E. coli* with a temperature sensitive defect in the native 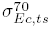 protein [37–40]. We first compare the structure of *Lactobacilli* sigma factors to other functional, heterologous sigma factors that have been expressed in *E. coli* in the past to detect any common features amongst these proteins. Two expression plasmids containing the *Lactobacilli* sigma factors promoter are then constructed and are shown to complement the defective 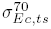 under non-permissive conditions. Potential applications for expression of foreign promoters in metabolic engineering are subsequently discussed.

## Experimental Procedures

### Bacterial strains and growth conditions

*E. coli* (BW25113 and 285c), *Lactobacillus brevis* (ATCC 367), and *Lactobacillus plantarum* (WCFS1) were used in this study. *E. coli* cultures were grown in LB broth at 37°C; 50 *µ*g/mL of streptomycin was added as necessary. The *E. coli* mutant strain 285c with a temperature sensitive defect in its *σ*^70^ protein and the parent strain P9OA5c [37, 41, 42] were generously provided by Robert Calendar (UC Berkeley) and cultured at 30°C in LB media and 25 *µ*g/ml streptomycin (if needed) to prevent selection of revertant mutations in *σ*^*H*^ [40]. For the complementation assay, the *E. coli* 285c strain were transformed with empty pCLE vector backbone or plasmids containing *L. brevis* or *L. plantarum σ*-factors. Cultures were grown overnight at 30°C in LB media with 25 *µ*g/ml streptomycin. The growth of the P9OA5c, 285c/pCLE, 285c/pCLE-Lp, and 285c/pCLE-Lb strains was monitored spectrophotomerically at 42°C using a TECAN plate reader to assess the complementation effect of the heterologous sigma factors. Batch studies of growth at 43.5-46°C were conducted in a shaking water bath at 200 rpm.

### *σ*-factor selection

Putative *σ*^70^ growth factors were identified on the basis of sequence annotation (*L. plantarum*: LP-1926, *L. brevis*: LVIS-0756) [43] for study. Heterologous sigma factors were placed under the control of *P*_*lac*_ with the native 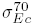 ribosome binding site to ensure equivalent translation efficiency. Primers for these sequences were designed from the *E. coli*-MG1655, *L. brevis* ATCC 367, and *L. plantarum* WCFS1 strains using Primer-BLAST [43] (Table S1). Sequence alignments of *E. coli*, *Lactobacilli*, and other sigma factor proteins [28–31, 44] using Clustal Omega [45].

### Genetic manipulation

The low copy number pCL1920 plasmid [46] was used to express the *σ*^70^-factors from *L. plantarum* and *L. brevis* in *E. coli*. First, the 300 base pair region upstream of the native *E. coli σ*^70^ gene containing the native RBS was introduced into pCL1920 to create pCL1920-EcS70 (pCLE). Note that the pCLE plasmid does not contain the 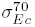 gene. Subsequently, each of the *σ* factor gene was amplified directly from the corresponding genomic DNA and subsequently purified using the Clean and Concentrator kit (Zymo Research). The fragments were digested with the appropriate restriction enzymes and ligated into the digested pCLE plasmid using T4 DNA ligase (New England Biolabs) to create 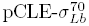 (pCLE-Lb) and 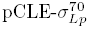 (pCLE-Lp). These expression plasmids were then introduced into *E. coli* (BW25113 and 285c) by electroporation (BioRad GenePulser XL) and plated onto selective media. All cloning PCR steps were performed using the Phusion polymerase (Finnzymes) and the constructs verified by sequencing (MCLAB Inc).

## Results

### Sequence Conservation in Heterologous Sigma Factors

In order to better understand why the Lactobacilli sigma factors are functional, we compared the sequences of all housekeeping sigma factors that have been expressed in *E. coli* to this date to identify any conserved structural features (Figure 3). All aligned sigma factors shared at least 44% amino acid similarity, and the *L. plantarum* and *L. brevis* factors are 66%-70% similar. Based on sequence alignments, domain 1 and the linker connecting it to domain 2 [13] appear to be entirely absent from the gram positive sigma factors, though there is a high degree of conservation in domains 2 and 4 in all of the heterologously expressed proteins. These sigma factor domains control interactions with RNA polymerase and DNA, and are therefore critical for promoter recognition [13]. Housekeeping sigma factors from *Caulobacter crescentus*, *Rickettsia prowazekii*, *Microcystis aeruginosa*, and *Bordetella pertussi* were also very similar to *E. coli σ*^70^ in domains 2 and 4, and all but *M. aeruginosa* contained a poorly conserved domain 1. Based on the data today, it appears that the only requirement for functional housekeeping sigma factor in *E. coli* may be sufficient similarity between domains 2 and 4 in the native and heterologous proteins [19].

### Lactobacilli sigma factors complement native *σ*^70^ defects

The first step in evaluating the properties of the heterologous sigma factors is to assess whether the sigma factors from *L. brevis* and *L. plantarum* are capable of complementing a temperature sensitive defect in the 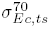 protein found in the *E. coli* 285c strain [37], as has been done repeatedly in the past for other heterologous sigma factors [29–31, 44, 47, 48]. First, low-copy plasmids containing the *Lactobacillus* sigma factors were introduced into *E. coli* 285c (see Methods and Materials). The growth kinetics of the 285c/pCLE-Lb, 285c/pCLE-Lp, 285c/pCLE, and the parent strains (P9OA5c) as shown in Figure 1 are quite similar at the permissive temperature (30°C), indicating that there is no defect in 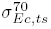 activity at low temperatures. However, subjecting the strains to the non-permissive temperature (42°C) reveals significant differences between the strains (Figure 2). Little to no growth is seen in the 285c/pCLE strain, while the 285c/pCLE-Lb and 285c/pCLE-Lp strains containing the heterologous sigma factors are not only viable, but indeed grow fairly robustly. The doubling time of the 285c/pCLE-Lb and 285c/pCLE-Lp strains are significantly greater than that of the parent strain (∼255 min versus 108 min for P9OA5c), suggesting that while 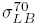 and 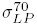 can restore viability at elevated temperatures, the growth rate of 285c is still impaired overall due to imperfect complementation or differences in promoter specificity compared to the native *σ*^70^. These results strongly support the conclusion that the *L. brevis* and *L. plantarum* sigma factors can successfully bind to the *E. coli* RNA polymerase and initiate transcription *in vivo* effectively, as otherwise the elevated temperature would have permanently arrested growth in all strains.

**Figure 1:**
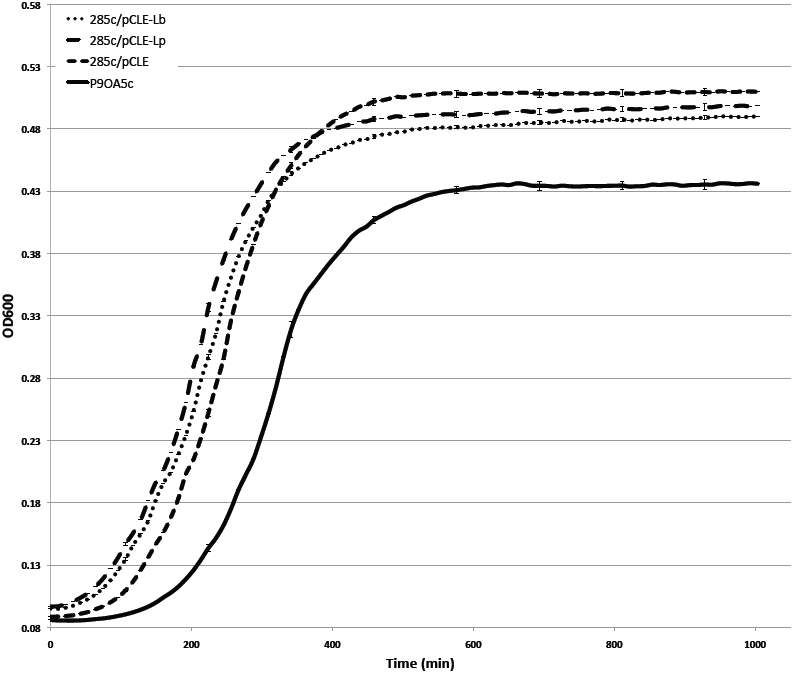
Growth of all 285c strains and the parent strain at 30°C. Each strain was inoculated from overnights into LB with 25 *µ*g/ml streptomycin (antibiotic omitted for P9OA5c) and cultured at 30°C to determine the growth kinetics at the permissive temperature. As can be seen, the growth rates and the yields of the strains are similar.

**Figure 2:**
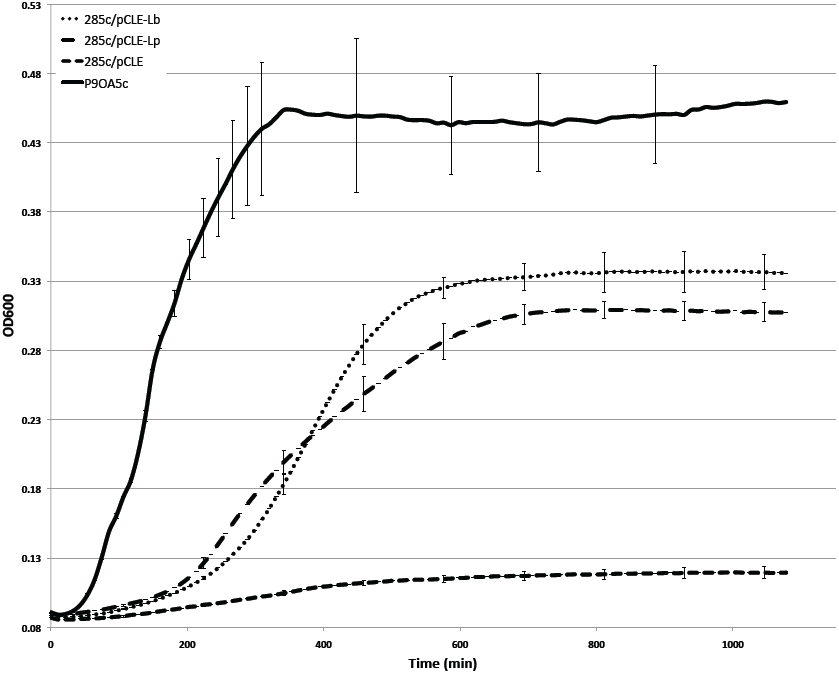
Complementation challenge at 42°C. Each strain was inoculated from overnights into LB with 25 *µ*g/ml streptomycin (antibiotic omitted for P9OA5c) and cultured at 42°C to determine if the heterologous sigma factors could complement a temperature sensitive defect in the native *σ*^70^ protein. Robust growth is observed for 285c/pCLE-Lb and 285c/pCLE-Lp while the 285c/pCLE strain is not viable at the elevated temperature.

**Figure 3:**
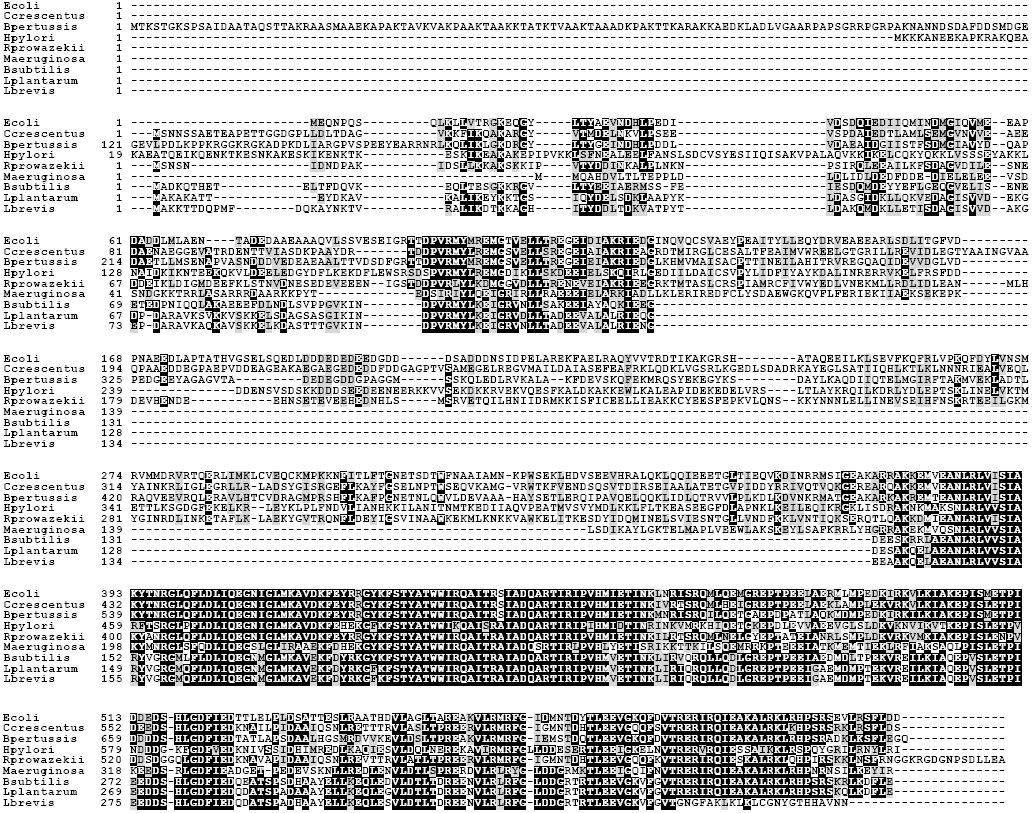
Alignments of *E. coli σ*^70^ with heterologously expressed *rpoD* homologs from other organisms [27–31, 44]. Conserved residues are shaded in gray or black depending while Alignments were performed with Clustal Omega [45] and visualized using Boxshade.

There are several possible physiological roles for the heterologous sigma factors within *E. coli* that can be inferred from this complementation effect. Firstly, these proteins may in fact act as direct replacements for 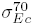 factor, enabling the transcription of genes required for exponential growth at high temperatures. This hypothesis is lent support by the fact that the *σ*^70^ promoter consensus sequences for lactic acid bacteria including those of *L. brevis* and *L. plantarum* are close to that of *E. coli* [49–54]. Efficient transcription of several *E. coli* promoters by the *Lactobacillus acidophilus* RNA polymerase holoenzyme has also been previously identified [55], providing additional weight to this interpretation. Suppressor mutations, such as those in *rpoH* or proteases, increase the stability of 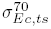 *in vivo* and restore the viability of 285c at elevated temperatures [56]. To test for compensatory mutations that restore growth at elevated temperatures, the ability of the 285c/pCLE-Lb and 285c/pCLE-Lp to grow at 43.5°C was assayed in batch cultures. Both strains grow robustly at 43.5°C (data not shown), indicating that they most likely do not have defects in proteolysis related genes that would stabilize 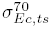. It is still possible that *rpoH* suppressor mutations exist in the strains but this phenotypic reversion is extremely rare (frequency ≈ 10^-8^) and unlikely to have occurred in independent replicates of 285c/pCLE-Lb and 285c/pCLE-Lp simultaneously. It was also recently demonstrated that expression of 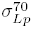 confers enhanced transcription of *L. planatarum* gene fragments [32], providing further evidence that these sigma factors are functional when heterologously expressed. Based on these growth kinetics and small likelihood of undetected compensatory mutations, we conclude that the *Lactobacillus σ*^70^ proteins are responsible for the observed complementation effect.

## Conclusions

We have demonstrated that two primary sigma factors from *L. brevis* and *L. plantarum* are able to complement a temperature sensitive growth defect in *E. coli* 285c that results from loss of the native *σ*^70^ activity at elevated temperatures. This finding appears to be one of the first demonstrating complementation with a Gram-positive primary sigma factor, and therefore provides additional insight into the plasticity of both the RNA polymerase core enzyme and the sigma factors themselves. Techniques that modify the native *E. coli* transcriptional program such as global transcription machinery engineering (gTME) may also see a benefit from the use of the *Lactobacilli* sigma factors given their ability to mediate transcription in complex with RNA polymerase. Engineering *E. coli* to transcribe *L. brevis* or *L. plantarum* genes effectively may also depend on the expression of the sigma factors in situations where compatible promoters cannot be introduced, such as during the protoplast fusion of these organisms.

## Acknowledgements

We gratefully acknowledge funding from the National Science Foundation Graduate Student Fellowship program, US NSF grant CBET-1032487, and the Texas Experimental Engineering Station, as well as experimental assistance from Maria Priscilla Almario Falls.

